# Single-Nuclear RNA Sequencing Reveals Regional Specialization and Cellular Interactions in Epicardial and Perivascular Adipose Tissue

**DOI:** 10.64898/2026.08.13.744748

**Authors:** Khanh-Van Tran, Bernard Ofosuhene, Anton Gulko, Taylor Orwig, Zinger Yang Loureiro, Christopher Jacobs, Bella Vogt, Irina Radu, David Bunsick, Linus Tsai, Leora Balsam, Jennifer Walker, Kate Fitzgerald, David McManus, Silvia Corvera, Evan D. Rosen, Margo P. Emont

**Affiliations:** Division of Cardiovascular Medicine, University of Massachusetts Medical School, Worcester, MA, USA; Division of Endocrinology, Diabetes, and Metabolism, Beth Israel Deaconess Medical Center, Boston, MA, USA; Division of Cardiac Surgery, University of Massachusetts Medical School, Worcester, MA, USA; Department of Medicine, University of Massachusetts Medical School, Worcester, MA, USA; Program in Molecular Medicine, University of Massachusetts Medical School, Worcester, MA, USA; Harvard Medical School, Boston, MA, USA; Broad Institute of MIT and Harvard, Cambridge, MA, USA; Section of Endocrinology, Diabetes, and Metabolism, Department of Medicine, University of Chicago, Chicago, IL, USA

## Abstract

**Background:** Adipose tissue surrounding the heart and vasculature plays critical roles in cardiovascular homeostasis and disease, yet the cellular and molecular milieu of these depots at single-cell resolution remains incompletely characterized. Understanding how regional adipocytes differ transcriptionally and communicate with neighboring cardiovascular cells is essential for developing targeted therapeutic strategies.

**Methods:** We performed single-nucleus RNA sequencing (snRNA-seq) on human adipose tissue from four anatomically distinct depots: ascending aorta, left atrium, right coronary artery, and subcutaneous fat. We characterized cellular composition, adipocyte and progenitor heterogeneity, depot-specific transcriptional programs, and intercellular communication networks. We further examined signaling remodeling in disease contexts, including atrial fibrillation and aortic aneurysm.

**Results:** We identified six transcriptionally distinct adipocyte subpopulations and six adipocyte stromal and progenitor cell (ASPC) subpopulations were shared across depots but showed marked differences in abundance and gene expression reflecting developmental imprinting, including HOX family genes and anterior–posterior patterning programs. Intercellular communication analysis revealed depot-specific ligand-receptor interactions, with EPHA signaling identified as selectively enriched in the left atrial adipose depot. Disease-state analyses demonstrated extensive change in cell-cell communication in atrial fibrillation and aortic aneurysm, with differential regulation of FN1, EGF, SLIT, NOTCH, and CD46 signaling pathways.

**Conclusions:** Our study reveals that cardiac and vascular adipose depots harbor transcriptionally specialized adipocytes and progenitors with distinct intercellular communication programs that are remodeled in atrial fibrillation and aortic aneurysm.

## Introduction

Adipose tissue is a metabolically dynamic organ system with depot-specific roles in energy storage, thermoregulation, and immune signaling^1^. Among its regional variants, epicardial adipose tissue (EAT) and perivascular adipose tissue (PVAT) have emerged as key players in cardiovascular physiology and pathology due to its anatomic proximity to the myocardium and coronary vasculature^2^. EAT particularly is thought to exert direct paracrine and vasocrine effects, as it is uniquely situated between the pericardium and myocardium, allowing for direct contact with cardiomyocytes^2^.

Clinical imaging studies have demonstrated that EAT volume and density, assessed by computed tomography, independently predict coronary artery disease burden, plaque vulnerability, and incident cardiovascular events^3,4^. Similarly, pericoronary adipose tissue attenuation on CT reflects local inflammatory activity and predicts myocardial infarction risk^5^. Despite these clinical associations, the cellular heterogeneity and transcriptional differences of EAT and PVAT remain incompletely characterized^6^. While prior studies have individually characterized human PVAT at the single cell level^7–9^, whole cell single cell studies cannot capture mature adipocytes due to the size and fragility of mature adipocytes^10^. Studies have begun to emerge that characterize human EAT at the single nuclear level^11^, capturing mature adipocytes, but none have characterized multiple EAT and PVAT depots at the single nuclear level and across the same individuals, limiting the ability to control for inter-individual variability in age, sex, comorbidities, and systemic metabolic status.

To address these gaps, we applied single-nucleus RNA sequencing (snRNA-seq) to profile the cell types present in EAT, PVAT and subcutaneous adipose tissue (SAT) within the same individuals undergoing cardiac surgery. By performing within-subject comparisons across the Left Atrium (LA), Right Coronary Artery (RCA), and Ascending Aorta (AA) depots to SAT from the anterior chest wall, we were able to control for confounding interindividual variability and gain a clearer picture of depot-specific transcriptional programs. In addition, to better understand the role of adipose tissue in cardiovascular disease, we analyze how disease states alter intercellular communication between adipose tissue and adjacent cardiac and vascular cells. Specifically, we focus on atrial fibrillation and aortic aneurysm as representative disease models of myocardial and vascular pathology. By examining disease-associated changes in signaling interactions, we aim to better understand how EAT and PVAT may participate in the pathological development of atrial fibrillation and aortic aneurysm.

## Methods

### Study Population and Sample Collection

Participants undergoing coronary artery bypass grafting (CABG), aortic valve replacement (AVR), or mitral valve repair/replacement (MVR) were prospectively enrolled at UMass Chan Medical School/UMass Memorial Medical Center in 2023–2025. Adipose tissue was collected from four anatomical regions: the right coronary artery (RCA), left atrium (LA), ascending aorta (AA), and subcutaneous adipose tissue (SAT) from the anterior chest wall at the inferior aspect of the midline incision. During clinically indicated surgery, approximately 40–100 mg of tissue was biopsied from each site under direct visualization by the cardiothoracic surgeon. Fat was immediately frozen in liquid nitrogen and sent for snRNA-seq. All samples were de-identified and stored at –80°C. This study was approved by the UMass Chan Medical School Institutional Review Board, protocol Study ID 000174. All subjects gave written informed consent. Samples and clinical data were stored under IRB-approved protocols and coded with a unique subject ID. Only authorized personnel had access to identifiable data.

### Nuclear Isolation and Single-Nuclear RNA Sequencing

Adipose samples were prepared and sequenced as previously described^12^. Briefly, samples were dissociated in Tween with salt and Tris (TST) buffer using a gentleMACS Dissociator (Miltenyi Biotec), filtered through 40 µm and 20 µm nylon filters (CellTreat), and incubated with individual hashtag antibodies for each sample (BioLegend) before flow sorting using a Beckman Coulter MoFlo Astrios EQ with a 70 μm nozzle to remove poor quality nuclei and count the same number of nuclei per sample. Samples were loaded onto a 10x Genomics Chromium Controller (10x Genomics) according to the manufacturer’s protocol. Single Cell 3’ v3.1 chemistry was used to process all samples and cDNA and gene expression libraries were generated according to the manufacturer’s instructions (10x Genomics). Gene expression libraries were multiplexed and sequenced on the Nextseq 500 (Illumina).

### Data Processing and Analysis

Raw sequencing data were processed using the Cell Ranger pipeline (10x Genomics, version 7.1.0) using GENCODE annotation GRCh38. DemuxEM^13^ (version 0.1.6) was used to sort cells into individual samples based on their antibody hashtags. Ambient RNA was removed from the processed DGEs using cellbender^14^ (version 0.2.0), and doublet scores were calculated using both scDblFinder^15^ (version 1.2.0) and scds^16^ (version 1.14.0). Cells were removed as doublets if they were both determined to be a doublet using scDblFinder and if they had a scds hybrid score > 1.5. Cells with < 800 UMIs were removed from the dataset, as were cells with > 10% of mitochondrial reads. Genes found in fewer than 2 cells were similarly removed. The data were then normalized using SCTransform and integrated using RPCA in Seurat^17^ (version 5.1.0), integrating by 10x run so as not to integrate over differences seen between depots. To subcluster, cells were subset into broad cell types and reintegrated and clusters were recalculated. Subclusters with high doublet scores and/or high percentages of mitochondrial reads were removed and the data was again reintegrated and clusters were recalculated. When the quality control metrics RNA count, feature count, mitochondrial percent, and scDblFinder score were plotted on calculated clusters and subclusters, none appeared to be strong outliers based on these metrics (**Supplementary Figures 1-2**) Cells were annotated by reference mapping^17^ to our previously published human adipose snRNA-seq data^18^ and by manually searching marker genes of clusters not present in the reference dataset.

### Pseudobulk and pathway analysis

To calculate pseudobulk counts, objects for individual cell types (adipocytes, or ASPCs) were split by sample and counts for each gene were summed across samples. Differential expression analysis (DEA) was run on pseudobulk samples using edgeR^19^ (version 4.2.2). UpSet plots were generated using UpSetR^20^ (version 1.4.0). Pathway analysis was performed with clusterProfiler^21^ (version 4.12.6) using GO biological processes pathways.

### Prediction of cell-cell interactions

To predict cell-cell interactions, data were subset to exclude “stromal_epithelial”, “neuronal” and “muscle” clusters due to low cell number, and split by depot. Datasets were then combined with either cardiomyocytes or vascular smooth muscle cells (VSMCs) from published datasets according to their location: LA EAT cells were combined with left atrial cardiomyocytes from healthy or AFib hearts^22^, RCA EAT cells were combined with right ventricle cardiomyocytes from healthy hearts^23^, and AA EAT cells were combined with aortic VSMCs from healthy aortas or aortic aneurysm^24^. To compare, SAT cells were also combined with healthy cells from all tested datasets. Each resulting dataset was analyzed using CellChat^25^ (2.1.2), filtering the output to exclude interacting clusters with fewer than 5 cells. Comparison analysis was performed between healthy samples and EAT versus SAT or between EAT and healthy versus unhealthy samples using CellChat. Venn diagrams were created using ggvenn^26^ (version 0.1.19) using pathways labeled as significantly upregulated in EAT cells compared to SAT cells using rankNet weight comparison in CellChat.

### Data Availability

Processed data are available to explore on the Broad Institute Single Cell Portal (<u>SCP3038</u>) and will be available to download from the Single Cell Portal and NCBI GEO upon publication. All raw sequencing data will be deposited in dbGaP in compliance with NIH data sharing policies. Other datasets used are the human adipose snRNA-seq^18^ (GSE176171), left atrium healthy and AFib snRNA-seq^22^ (GSE255992), ascending aorta healthy and aneurysm snRNA-seq^24^ (GSE207784), and healthy right ventricle snRNA-seq^23^ (ERP123138).

## Results

### A single cell atlas of epicardial and perivascular fat

We performed snRNA-seq on three epicardial/perivascular fat depots (LA, RCA, and AA) as well as paired SAT samples from 10 individuals (**Table 1**, **Figure 1a**). After data processing we were left with 23,207 total cells (4,781 AA, 6,946 LA, 6,205 RCA, and 5,275 SAT) (**Figure 1b**). The data were integrated by individual and thus the clustering showed little difference across individual, sex, or BMI range (**Supplementary Figure 1a-c**). While most cell types were found in all depots, mesothelial cells were present in the EAT/PVAT depots but not the SAT, as expected. Adipocytes were found to comprise approximately 50% of SAT cell types but only 25% or less of EAT/PVAT cells, with the remaining percentage made up by the aforementioned mesothelial cells as well as an increased proportion of vascular cells compared to SAT (**Figure 1c**). Interestingly, we also identified a population resembling neuronal cells selectively in the LA samples, consistent with prior findings that ganglionated plexi are found in epicardial fat^27,28^

**Figure 1.**
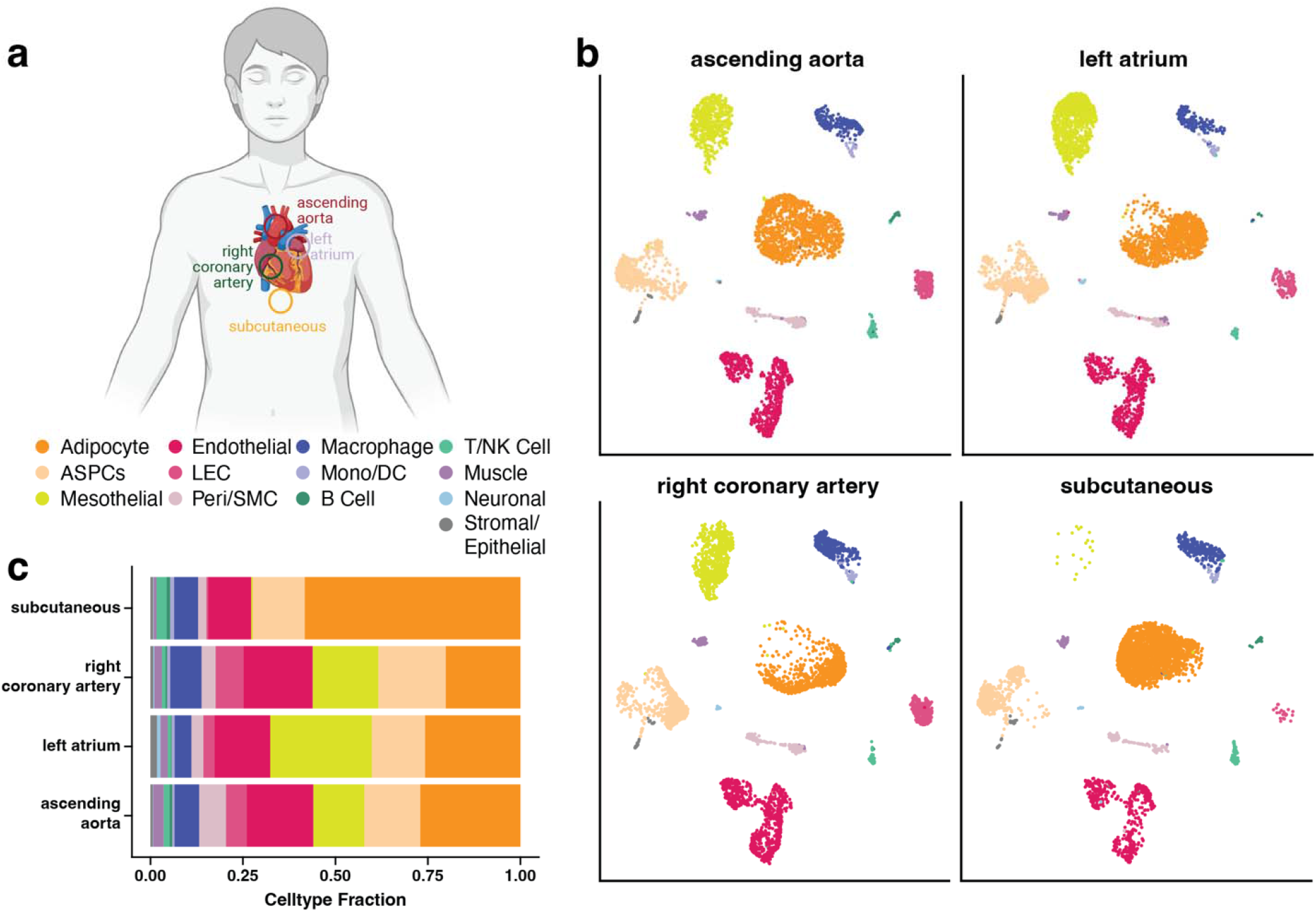
Single cell profiling of epicardial adipose tissue. **a,** Diagram showing locations of adipose tissue samples. **b,** UMAP plot of integrated snRNA-seq analysis of adipose tissue, split by depot and subsampled to the same number of cells (4781) per depot. **c,** Proportion of cell types in adipose samples split by depot.

**Table 1:**
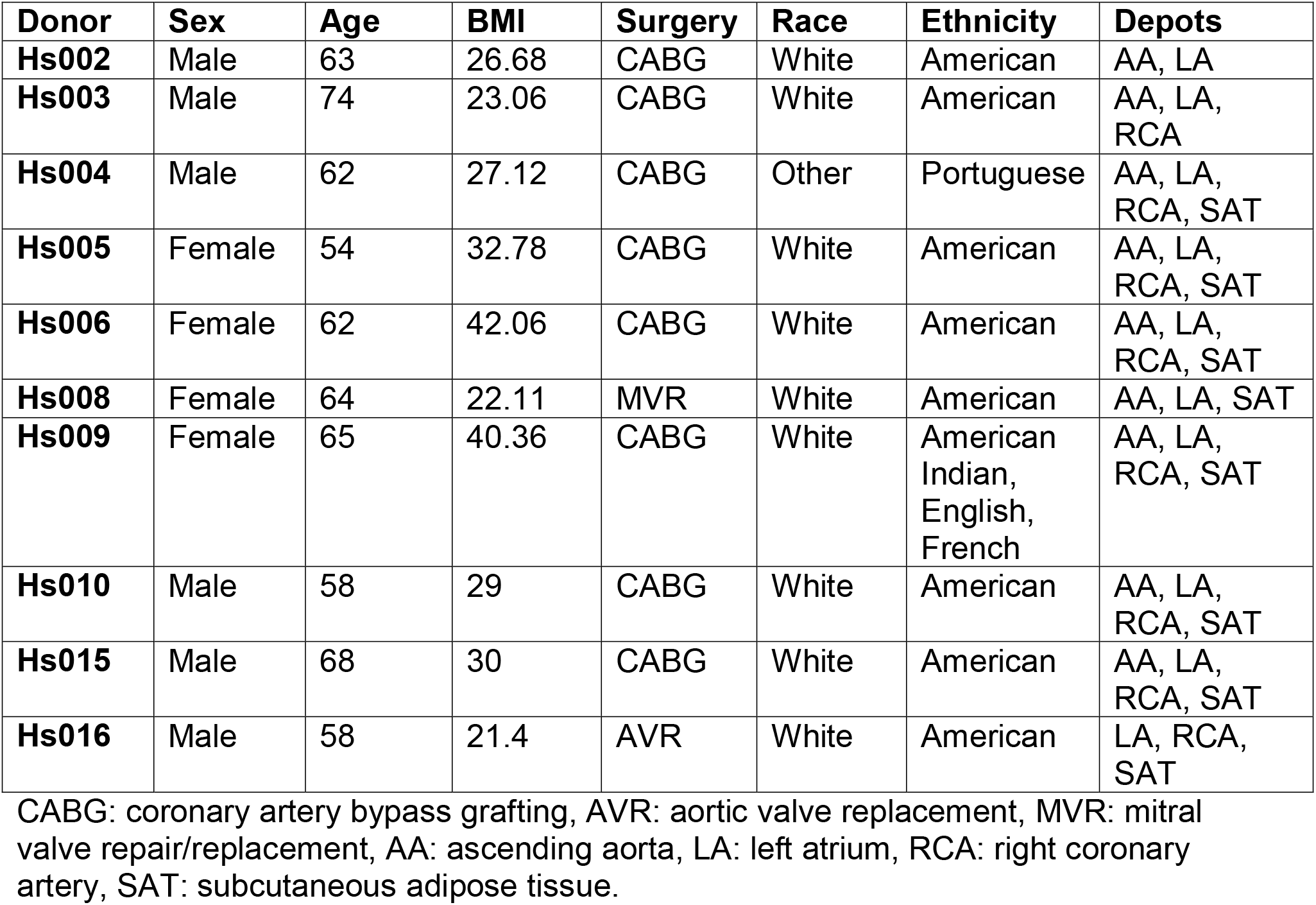
Subject information.

### Identification of Transcriptionally Distinct Adipocyte States Across Depots

While the proportions of mature adipocytes in the EAT/PVAT depots were roughly similar, we hypothesized that adipocyte gene expression might be different across depots, particularly as a consequence of proximity to the heart. We therefore subclustered the adipocytes from our integrated object, identifying six discrete transcriptional states (**Figure 2a&b**). While most subclusters were found across all depots, there was some depot selectivity, most notably Ad^KLF2^ was most abundant in SAT and AA samples, Ad^GATA4^ was only found in EAT/PVAT samples, and Ad^RGS6^ was most abundant in AA and LA samples (**Figure 2c**). Pathway analysis of the marker genes of these subclusters suggested that some adipocytes subtypes may have specialized functions, for example marker genes for the EAT/PVAT selective population Ad^GATA4^ were enriched for genes in the “Artery Development” pathway while the “Adaptive thermogenesis” pathway was enriched in Ad^RGS6^ marker genes (**Figure 2d&e**).

**Figure 2.**
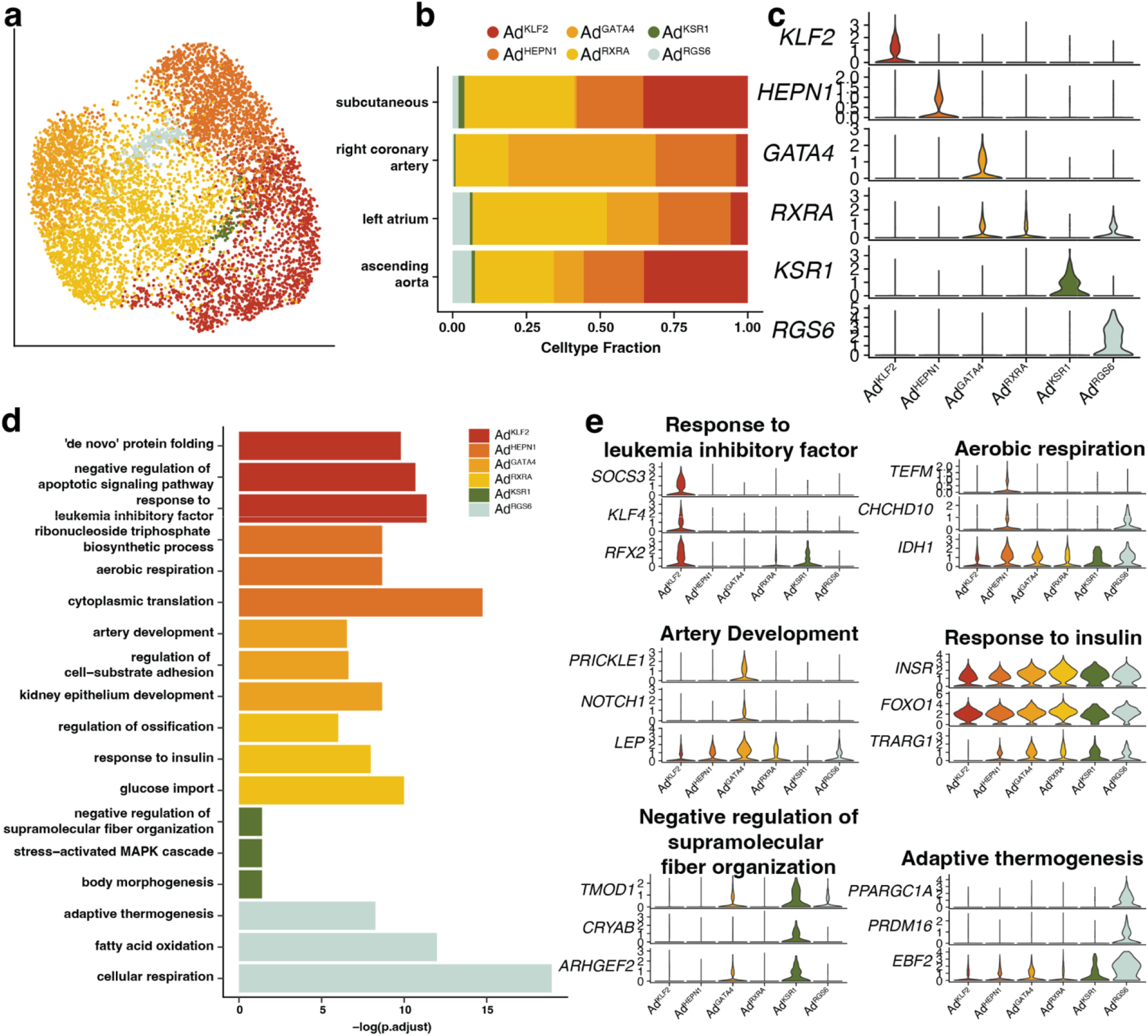
Adipocyte subclusters and gene expression vary across adipose depot. **a,** UMAP plot of integrated adipocyte subclusters (n = 7513 cells). **b,** Subcluster distribution across depot. **c,** Marker genes for adipocyte subclusters. **d,** Selected GO pathways from pathway analysis of adipocyte subcluster marker genes. **e,** Selected marker genes highlighted in pathway analysis.

We next examined the regulation of genes associated with common adipocyte processes; secretion of signaling molecules, response to insulin, lipid handling, and thermogenesis across adipocyte subcluster and depot (**Supplementary Figure 3a, b**). While the plotted genes were expressed in most adipocyte subclusters and depots, adipokines such as LEP and CFD appeared to be enriched in the two depots closest to the heart, the RCA and LA, compared to the AA and SAT. Conversely, lipogenesis genes such as DGAT1 and DGAT2 were higher in the SAT. Thermogenic genes, particularly those associated with classical thermogenesis were somewhat associated with LA adipocytes and, as suggested by the pathway analysis, more strongly associated with Ad^RGS6^, an LA and AA enriched subpopulation. Intriguingly, genes associated with both classical thermogenesis and creatine cycling were associated with this population.

To harmonize results with previous studies, we used reference mapping to compare the identified adipocyte populations to our previous work (**Supplementary Figure 3c**). While there was not strong convergence across all populations, we did see that the majority of cells in the Ad^KSR1^ population mapped to our previous hAd5 population, a population that was associated with response to signaling in our previous work, and that the majority of Ad^RGS6^ cells map to our previous hAd6 population, a visceral population that was also found to express thermogenic markers. Because adipocyte gene expression is sensitive to factors such as BMI and depot, the differences in patient population and sample depot likely explain the lack of clear mapping across the rest of the subpopulations.

### Stromal Progenitor Cells Exhibit Regional Specialization

We next sought to investigate the adipose stromal and progenitor cells (ASPCs) to determine if these too showed gene expression differences correlated with their proximity to the heart. Similarly to the adipocytes, we subclustered the ASPCs and found strong location-based separation in the calculated subclusters. Subpopulations ASPC^SOCS3^ and ASPC^FHOD3^ were largely found in the SAT and AA fat, while subpopulations ASPC^CNTNAP2^, ASPC^NR4A3^, and ASPC^CD81^ were more abundant in the RCA and LA (**Figure 3a-c**). When we performed pathway analysis on the marker genes of these subpopulations we found that ASPC^SOCS3^ was enriched in the “Positive regulation of miRNA transcription” pathway, and the specific marker genes in this pathway are largely adipocyte differentiation genes, while ASPC^FHOD3^, was associated with “Cell-matrix adhesion”. EAT-selective populations ASPC^CNTNAP2^, ASPC^NR4A3^, and ASPC^CD81^ were associated with “Cyclic-nucleotide mediated signaling”, “Cardiac muscle tissue development” and “Protein folding”, pointing to possible cardiac-selective processes in these cells (**Figure 3d&e**).

**Figure 3:**
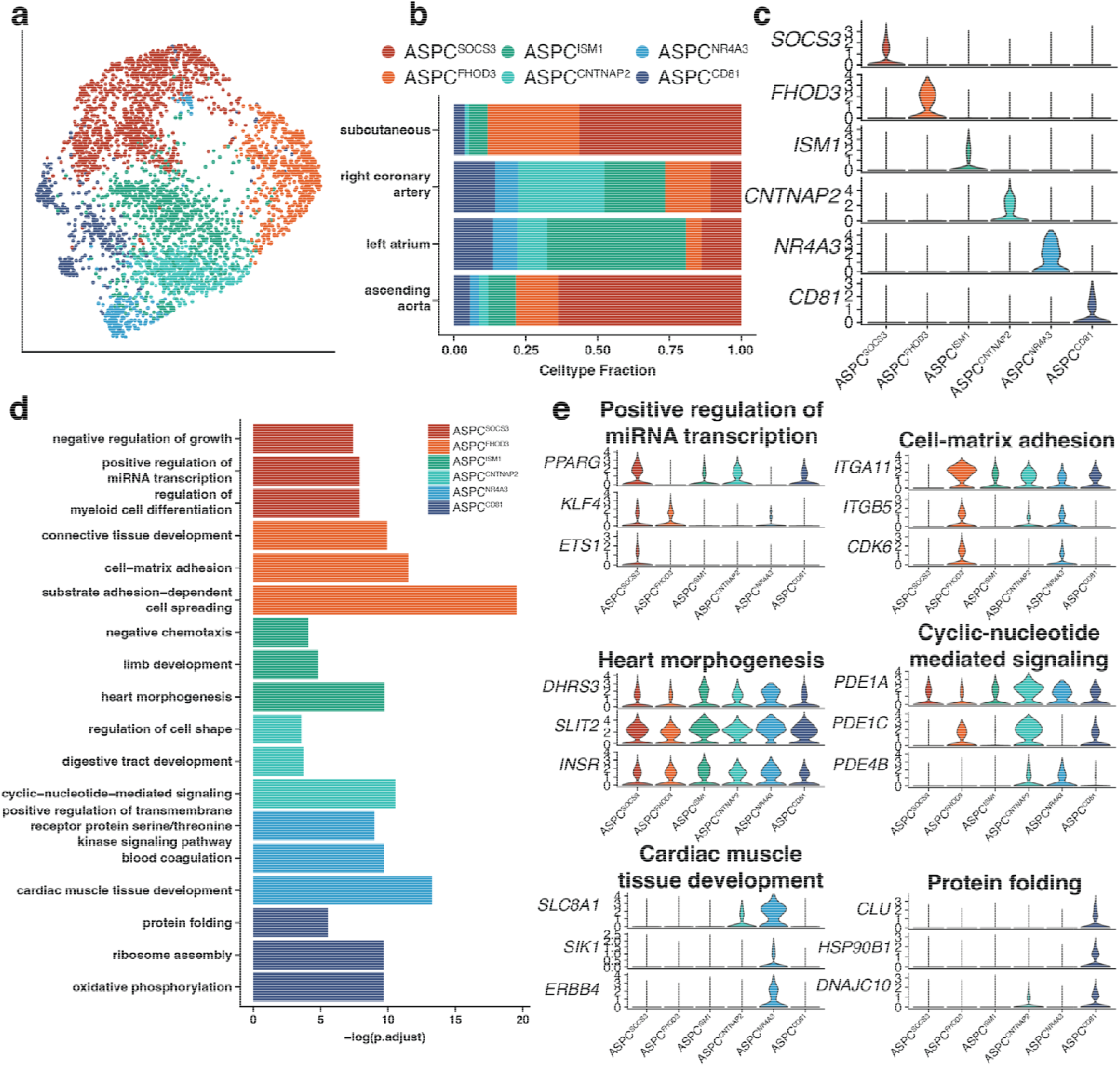
ASPC subclusters and gene expression vary across adipose depot. **a,** UMAP plot of integrated ASPC subclusters (n = 3593 cells). **b,** Subcluster distribution across depot. **c,** Marker genes for ASPC subclusters. **d,** Selected GO pathways from pathway analysis of ASPC subcluster marker genes. **e,** Selected marker genes highlighted in pathway analysis.

As with the adipocytes, we mapped our ASPC subpopulations to those from our previous work (**Supplementary Figure 4d**). 66% of ASPC^SOCS3^ cells mapped to our previously identified “committed preadipocyte” population hASPC1 and 84% of ASPC^FHOD3^ cells mapped to the DPP4+ “early ASC” population hASPC2, suggesting that these populations are similar. Additionally, most cells that mapped to the hASPC4 “Areg” population were in cluster ASPC^ISM1^. The cardiac-selective populations ASPC^CNTNAP2^, ASPC^NR4A3^, and ASPC^CD81^ did not map as strongly to any distinct previously identified subclusters.

### Depot-specific regulation of adipocytes and progenitor cells

To further investigate the overall differences between adipocytes and ASPCs in different depots, we performed pseudobulk analysis, in which we ran differential expression analysis on summed counts for a specific cell type across samples. A principal component analysis (PCA) plot of the adipocyte pseudobulk data finds clear separation between SAT and EAT adipocytes, with the strongest differences found between the SAT and RCA samples (**Figure 4a**). Differential expression analysis between each EAT/PVAT depot and SAT found differential genes across all depots, with the majority of EAT/PVAT up- or down-regulated genes being specific to LA or RCA or shared across both (**Figure 4b,c**). Pathway analysis on LA & RCA shared differential genes found that upregulated genes tended to be associated with growth pathways such as ‘embryonic organ morphogenesis’, while shared downregulated (and therefore SAT upregulated) genes were associated with positioning and developmental pathways like ‘anterior/posterior pattern specification’ (**Figure 4d**). Pseudobulk analysis of the ASPCs also found a separation between SAT and EAT/PVAT ASPCs, with the most differential genes found between the SAT and the RCA and LA (**Figure 4e-g**). Pathway analysis on the shared LA and RCA upregulated genes found that ‘response to BMP’ and ‘cardiac muscle tissue development’ were upregulated in EAT ASPCs, while ‘regulation of chemotaxis’ and ‘regulation of leukocyte migration’ were upregulated in the SAT ASPCs (**Figure 4h**).

**Figure 4.**
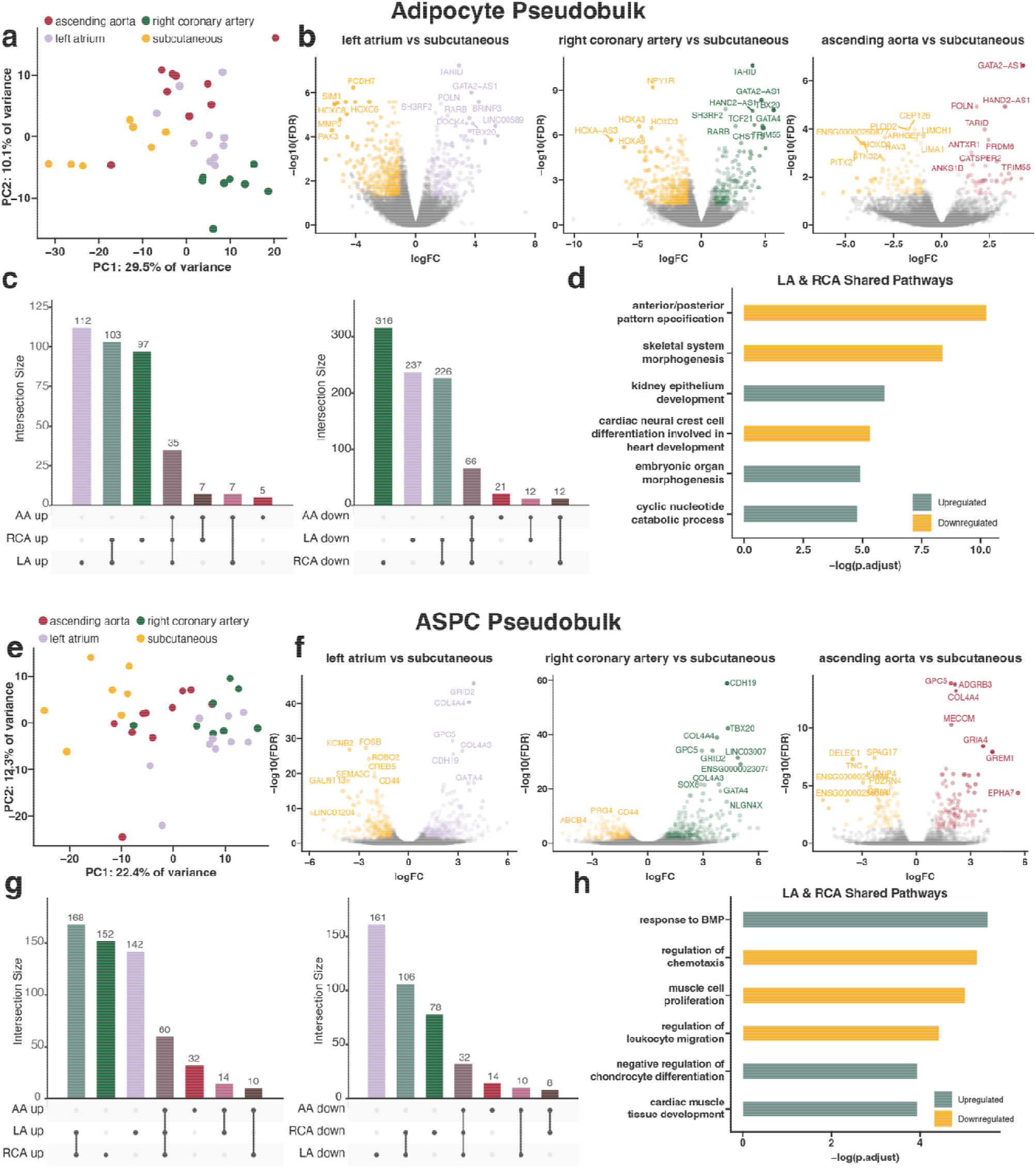
Pseudobulk analysis of adipocytes and ASPCs by depot. **a,** PCA analysis of adipocyte pseudobulk data. **b,** Volcano plots comparing gene expression in adipocytes from each epicardial depot compared to subcutaneous adipocytes. **c,** UpSet plot showing the intersection between upregulated (left) or downregulated (right) genes in epicardial adipocytes compared to subcutaneous adipocytes. **d,** Pathway analysis of genes upregulated (green) or downregulated (yellow) in both left atrium and right coronary artery adipocytes compared to subcutaneous adipocytes. **e,** PCA analysis of ASPC pseudobulk data. **f,** Volcano plots comparing gene expression in each ASPCs from epicardial depot compared to subcutaneous ASPCs. **g,** UpSet plot showing the intersection between upregulated (left) or downregulated (right) genes in epicardial ASPCs compared to subcutaneous ASPCs. **h,** Pathway analysis of genes upregulated (green) or downregulated (yellow) in both left atrium and right coronary artery ASPCs compared to subcutaneous ASPCs.

We next examined the regulation of HOX genes, which are responsible for developmental patterning and thus would be expected to be differentially expressed across adipose depots. When we plotted HOX genes specifically on the pseudobulk analysis of adipocytes and ASPCs we found a number of these genes were differentially expressed (**Supplementary Figure 4a&b**). Surprisingly, the expression of HOX genes was most differential in the adipocytes compared to the ASPCs, and significantly enriched in the subcutaneous but not epicardial tissue. Plotting these genes directly on the adipocyte single cell data found that HOX gene expression was highest in the Ad^KSR1^ subcluster and that again expression of these genes was most strongly found in subcutaneous adipocytes (**Supplementary Figure 4c**).

### Vascular, immune and mesothelial subcluster proportions shift across depot

To examine the other cell types in our dataset we further subclustered the vascular cells, immune cells, and mesothelial cells (**Supplementary Figure 5**). Vascular cells overall resembled those that we have described previously^18^. For ease of analysis we included the noted neuronal population as well as populations resembling cardiomyocytes or other muscle cells in this object; it is unknown if these cells are present in the EAT or if they are present due to carryover from the tissue isolation. While proportions of blood endothelial cells are similar across depots, lymphatic endothelial cells appear to be enriched proportionally in EAT/PVAT compared to SAT (**Supplementary Figure 5a-c**).

A relatively small number of immune cells were isolated from these samples compared to our previous studies, with immune cells comprising 6-12% of cells per studied depot. This is in contrast to our previous work^18^ in which immune cells comprised 17% of VAT cells and 24% of SAT cells, possibly due to the differences in patient characteristics between the two studies. Within these cells we were able to identify three populations of macrophages, one combined population of monocytes and dendritic cells, a population of B cells, and a population of T/NK cells (**Supplementary Figure 5d-f**). The RCA was enriched in macrophages compared to other depots, specifically in the increased proportion of Mac^ABCA9^ and the emergence of the Mac^AGBL4^ population which is largely found in this depot. The Mac^FOS^ population is conversely increased in the SAT and AA depots.

Mesothelial cells were found on the visceral EAT/PVAT samples, but not in the SAT samples, as expected (**Figure 1b,c**). Subclustering the mesothelial cells found 4 populations that were influenced by depot. Mes^CCND3^ was higher in the LA samples, while Mes^NFATC2^ was enriched in RCA and AA and Mes^EBF1^ was specifically enriched in AA (Supplementary **Figure 5g-i**).

### Prediction of cell-cell communication suggests regional differences

We next sought to examine the functional consequences of the observed regional differences by predicting cell-cell communication between EAT/PVAT cells and cardiomyocytes/vascular smooth muscle cells (VSMCs). To do this, we downloaded single nuclear datasets of the left atrium^22^, right ventricle^23^, and ascending aorta^24^, and subset healthy cardiomyocytes or VSMCs as applicable. We combined these cells with our datasets by depot, using the closest EAT/PVAT depot as well as SAT cells as a control (**Figure 5a**). We then used CellChat^25^ to predict interactions between all cells in the combined datasets, and examined the strength of predicted interactions in the EAT/PVAT dataset compared to SAT (**Figure 5b**). In all comparisons, the strength of predicted interactions between cardiomyocytes or VSMCs and the analyzed EAT/PVAT depot was higher for the majority of cell types compared to SAT, suggesting that some of the observed regional differences seen may be related to signaling with neighboring cardiac cells. We next specifically interrogated the similarity in interactions between cardiomyocytes/VSMCs and adipocytes/ASPCs that were significantly upregulated in EAT/PVAT compared to SAT (**Figure 5c**). We found that many of the significant interactions were shared, either between all three depots or specifically between LA and RCA. One unique interaction that stood out was EPHA, which is only upregulated in LA in 75% of the queried interactions. Specifically plotting the EPHA signaling pathway, we found increased predicted interactions between cardiomyocytes and ASPCs in the LA compared to the SAT, and plotting genes related to EPHA signaling in our dataset finds that *EPHA3* expression is found in all pericyte/SMCs, but is only found in LA ASPCs (**Figure 5d,e**), suggesting that LA adipose tissue has a unique capacity to respond to some cardiomyocyte signaling.

**Figure 5:**
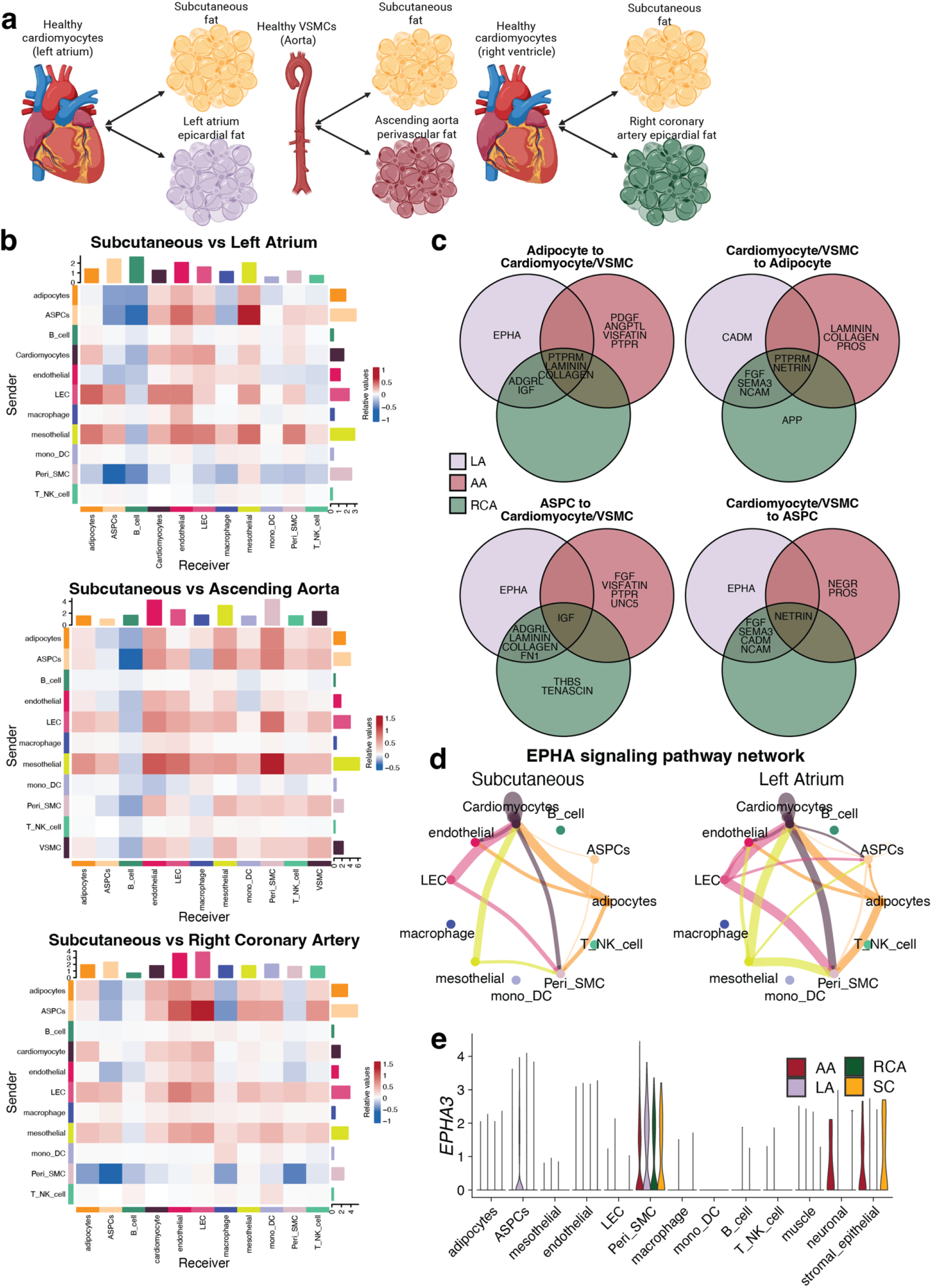
Cellchat analysis predicts epicardal fat-selective interactions with cardiomyocytes. **a,** Diagram describing the cellchat analysis experiments. Data from each epicardial depot was combined with cardiomyocyte or vascular smooth muscle cell data from a corresponding cardiac or vascular dataset and cellchat was run on this combined dataset. Subcutaneous data was similarly combined with each cardiac and vascular dataset to serve as a control. **b,** Heatmaps showing the predicted interaction strength in each epicardial dataset compared to subcutaneous. **c,** Comparison of predicted interactions that were significantly upregulated between epicardial and cardiovascular cell types compared to subcutaneous and cardiovascular cell types. **d,** Chord plot showing the predicted EPHA signaling pathway network in subcutaneous versus Left Atrium fat. **e,** Violin plot showing expression of the receptor gene *EPHA3* in all cell types, split by depot.

### Prediction of cell-cell communications suggests interaction with disease states

We next sought to use our data to predict the interactions between adipose tissue and cardiovascular cells in cardiovascular disease states. To do this we used the previously downloaded left atrium dataset, which included both healthy samples as well as those from atrial fibrillation (AFib). We combined the cardiomyocytes from healthy or AFib left atria with our left atrium epicardial fat data and again predicted cell-cell interactions (**Figure 6a**). We see that, unsurprisingly, predicted interactions between adipose cell types are similar across comparisons as in this analysis the adipose data remains the same. However, predicted interactions between adipose cell types and cardiomyocytes are almost uniformly increased in AFib cardiomyocytes compared to healthy ones (**Figure 6b**). When examining changes in the specific predicted interactions, we saw that FN1 signaling, which is found between cardiomyocytes and multiple adipose cell types in the healthy state, is no longer present in the disease state. Conversely, EGF signaling is not found between cardiomyocytes and adipose cell types in the healthy state but emerges in the AFib cardiomyocytes (**Figure 6c**). This suggests that cardiovascular disease is associated with changes in cell signaling between cardiomyocytes and adipose tissue.

**Figure 6.**
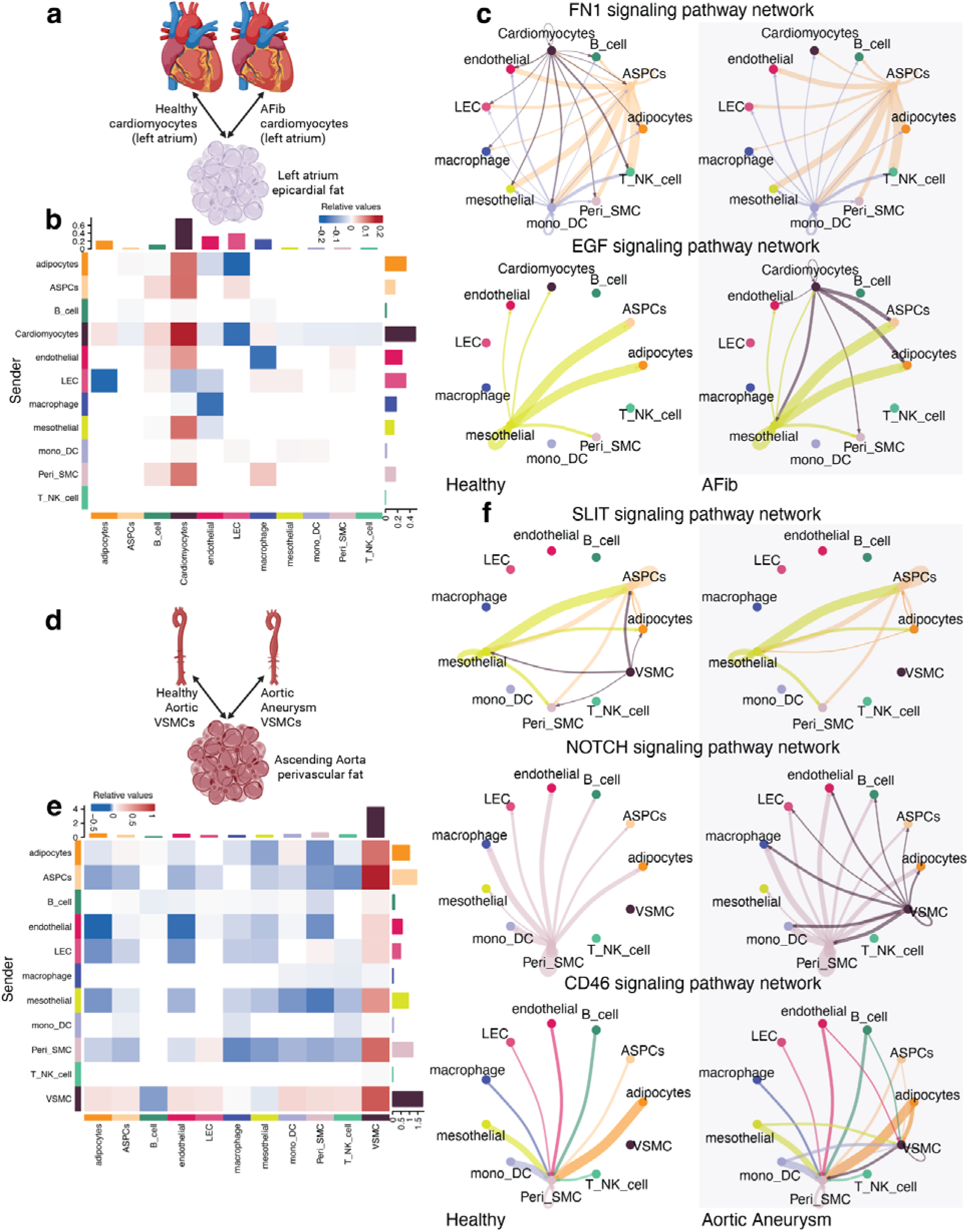
Cellchat analysis predicts changing interactions between fat and cardiovascular cells in the disease state. **a,** Diagram of performed cellchat analysis. Single nuclear data from left atrium fat was combined with cardiomyocytes from either healthy or AFib left atrium hearts, then predicted interactions were compared across disease state. **b,** Heatmap showing the strength of predicted interactions with AFib versus healthy cardiomyocytes. **c,** Select signaling networks predicted between left atrium epicardial fat and healthy versus AFib cardiomyocytes. **d,** Diagram of performed cellchat analysis. Single nuclear data from ascending aorta fat was combined with VSMCs from healthy aorta versus aortic aneurysm. **e,** Heatmap of strength of predicted interactions in aortic aneurysm compared to healthy aorta. **f,** Selected signaling networks predicted between ascending aorta epicardial fat and VSMCs from healthy or aneurysm aortas.

To predict the effects of disease on VSMC/adipose tissue interactions we next returned to the aorta dataset, which included data from healthy samples as well as from aortic aneurysm. Again, we combined the VSMCs from both conditions with our AA PVAT sample and predicted cell-cell interactions (**Figure 6d**). As with AFib, the weight of the predicted interactions between VSMCs and adipose tissue cells was strongly increased in the aneurysm samples compared to control (**Figure 6e**). When looking at specific differential pathways we found multiple patterns, in the SLIT signaling pathway VSMCs signaled to multiple adipose tissue cells in the healthy state, but this signaling was lost in the disease state. Conversely with NOTCH signaling, VSMC signaling was activated in the disease state. For CD46 signaling, VSMCs didn’t participate in signaling at all in the healthy state but in the disease state they were now predicted to both send and receive signals.

## Discussion

This study provides a comprehensive single-nucleus transcriptomic atlas of human adipose tissue across four anatomically distinct depots, including AA, LA, RCA, and SAT. Our study demonstrates that while the major cellular constituents of adipose tissue are conserved across depots, both adipocytes and their progenitors segregate into transcriptionally distinct subpopulations with depot-specific patterns and differential transcriptional programs including HOX family genes. Moreover, intercellular communication networks differ markedly between cardiac/vascular and subcutaneous depots, with EPHA signaling enriched in LA adipose depot. These communication networks undergo significantly remodeling in atrial fibrillation and aortic aneurysm, implicating specific signaling pathways, including FN1 and EGF in atrial fibrillation and SLIT, NOTCH, and CD46 in aortic aneurysm.

### Regional Distinct Adipocyte States Across Depots

Our finding that cardiac and vascular adipocytes exhibit downregulation of HOX genes and anterior-posterior patterning signaling pathways compared to subcutaneous adipocytes extends the concept of developmental imprinting from the established subcutaneous-visceral axis to the far less explored cardiac depot axis^29,30^. Prior work by Karastergiou et al. demonstrated that visceral and subcutaneous adipocytes express distinct HOX gene profiles that persist through in vitro culture, indicating cell-autonomous epigenetic regulation^29^. Our results indicate that HOX gene expression is less prominent in epicardial and perivascular adipocytes, suggesting that EAT and PVAT possess a developmental transcriptional program distinct from that of subcutaneous adipocytes. Beyond this developmental signature, we find adipocyte populations exhibit marked subtype heterogeneity. Our findings suggest that LA and AA may have higher thermogenic competent cell populations (Ad^RGS6^), though the majority of cells do not have significant expression of UCP1 or other classical thermogenic markers, suggesting that EAT and PVAT most resemble visceral white adipose depots with thermogenic cells rather than a brown adipose depot.

We further observed depot-selective adipokine expression, with LEP and CFD preferentially expressed in RCA and LA adipocytes relative to AA and SAT, consistent with regional differences in the regulation of local metabolic and inflammatory homeostasis. Conversely, lipogenesis-associated genes including DGAT1 and DGAT2 were enriched in SAT, consistent with the established role of subcutaneous fat as the primary site of triglyceride storage and esterification.

### Regional Adipose Stem Progenitor Cell Identity

Our analysis revealed six transcriptionally distinct ASPC subclusters whose distribution varied substantially by anatomical location, with SAT and AA fat enriched for ASPC^SOCS3^ and ASPC^FHOD3^ and epicardial depots harboring higher proportions of ASPC^ISM1^, ASPC^CNTNAP2^, ASPC^NR4A3^, and ASPC^CD81^, suggesting that the tissue microenvironment shapes progenitor identity^31^. Differential expression analysis showed that epicardial ASPCs shared a common set of upregulated genes including GPC5, COL4A3/4, GATA4, and CDH19. Pathway analysis of genes differentially expressed in LA and RCA ASPCs highlighted upregulation of BMP signaling, negative regulation of chondrocyte differentiation, and dysregulation of cytoskeletal migration. Notably, immune-related pathways were significantly downregulated in cardiac relative to SAT ASPCs, consistent with the role of subcutaneous fat as a barrier tissue requiring robust immune surveillance against infection, whereas excessive inflammation in the cardiac compartment would be detrimental to myocardial function. EAT has upregulation of the BMP pathway, which is a well-established driver of adipogenesis and brown/beige adipocyte differentiation^32,33^. BMP signaling upregulation in epicardial ASPCs may be mechanistically linked to the thermogenic phenotype observed in Ad^RGS6^, raising the possibility of a BMP responsive progenitor-to-thermogenic-adipocyte axis specific to the epicardial environment. Pathways involved in the negative regulation of chondrocyte differentiation and cardiac muscle development are also upregulated in EAT, suggesting that EAT actively contributes to cardiac remodeling, perhaps, by suppressing these processes. Together, these findings demonstrate that human ASPCs are a functionally stratified compartment whose transcriptional programs reflect depot-specific roles.

### Depot-Specific Cellular Composition

Beyond adipocytes and their progenitors, several cell-type-level differences across depots were evident. Mesothelial cells were restricted to the visceral EAT/PVAT depots and absent from SAT, consistent with the mesothelial lining of the pericardial and periaortic space, and resolved into four depot-enriched subpopulations (Mes^CCND3^ enriched in LA, Mes^NFATC2^ in RCA and AA, and Mes^EBF1^ specifically in AA). A neuronal-like population was identified selectively in the LA, its low cell number warrants cautious interpretation and may reflect capture of signals from intrinsic cardiac autonomic ganglia embedded within the adipose tissue. Lymphatic endothelial cells were proportionally enriched in EAT/PVAT relative to SAT, potentially reflecting a role for cardiac adipose tissue in pericardial lymphatic drainage. Finally, the RCA depot was enriched in Mac ^ABCA9^ and Mac^AGBL4^ macrophage subpopulations relative to the Mac^FOS^-predominant SAT and AA; given the established link between pericoronary adipose tissue inflammation and coronary atherosclerosis burden and plaque vulnerability, this distinct macrophage composition may reflect an inflammatory microenvironment shaped by proximity to the coronary vessel wall^5^. The functional roles of these subpopulations in coronary pathophysiology require further investigation.

### Depot-Specific Adipose Signaling

Our cell-cell communication analysis reveals that regional differences in EAT and PVAT transcriptional identity are not merely intrinsic to the adipose compartment but are reflected in the predicted signaling relationships with neighboring cardiac and vascular cells. Across all depot-cardiac cell comparisons, interaction strength between EAT/PVAT and cardiomyocytes or VSMCs exceeded that of SAT, supporting the hypothesis that anatomic proximity, combined with the absence of fascial barriers separating EAT/PVAT from adjacent cardiovascular tissue, enables direct paracrine communication to modulate myocardial and vascular function^34,35^.

Among the shared and depot-enriched interactions, the most distinctive finding was the LA-specific upregulation of EPHA signaling. The ephrin receptor signaling is increasingly recognized as a regulator of tissue patterning, cell migration, and progenitor fate, though its role in cardiac adipose biology is less known^36^. Cardiomyocyte-ASPC interactions via EPHA signaling were specifically enriched in the LA compared to SAT as well as RCA and AA, this raises the possibility that LA cardiomyocytes actively shape the progenitor niche within the overlying adipose tissue in a depot-specific manner. Whether this reflects developmental programming of the LA EAT compartment or an adaptive response to the unique hemodynamic and electrophysiological environment of the left atrium remains to be determined.

### EAT and PVAT Signaling in Disease States

#### Atrial fibrillation

The increased number of predicted interactions between cardiomyocytes and LA EAT cells in AFib compared to healthy controls suggests active participation of adipose tissue in the atrial fibrillation^37^. Rather, AF-remodeled cardiomyocytes have significantly altered signaling programs. Because the adipose data were identical in both conditions, our findings reveal a shift in signaling pattern in the cardiomyocytes of AF patients. FN1 signaling from cardiomyocytes to adipose cells was lost in AFib, while EGF signaling emerged. Fibronectin is generally increased in the fibrillating atrium^38^, but that rise is fibroblast-driven^39^. Because our fibroblast-like cells (ASPCs) were held constant, our analysis isolates the cardiomyocyte contribution, which decreased. Given that these interactions are computational predictions, we regard this FN1 finding as hypothesis-generating and speculate that cardiomyocytes reduce FN1 signaling as fibroblasts assume the dominant role in matrix production. Conversely, the emergence of EGF signaling in AFib is consistent with activation of proliferative and pro-fibrotic programs that have been implicated in atrial remodeling^40^. These findings suggest that the transition from health to disease is accompanied not simply by an increase in adipose-cardiomyocyte signaling but by a qualitative shift in the nature of that communication.

#### Aortic Aneurysm

Our observation that the aggregate weight of predicted VSMC-adipose interactions is broadly increased in aneurysm versus control aorta is consistent with a model in which perivascular fat becomes a more active signaling partner of the vessel wall during disease, and suggests that direct, contact- and ligand-dependent crosstalk, beyond bulk adipokine secretion, may contribute to the VSMC phenotypic instability that characterizes aneurysm formation. The loss of SLIT signaling from VSMCs in the aneurysmal state is notable because Slit2, acting through Robo receptors, behaves as a chemorepellent that restrains VSMC migration; recombinant Slit2-N inhibits PDGF-stimulated VSMC migration by suppressing Rac1 activation and lamellipodia formation, identifying Slit2 as a brake on smooth muscle motility^41^. Loss of this signal in disease could therefore be permissive for the migratory, dedifferentiated VSMC behavior seen early in aneurysm formation. Conversely, the emergence of NOTCH signaling aligns well with experimental work tying Notch activation to AAA: NICD and Hes1 are elevated in human and angiotensin II–induced murine AAA tissue, and Notch1 drives the VSMC phenotypic changes^42,43^. Finally, the appearance of CD46 signaling in the diseased aorta is intriguing given that CD46 is a membrane complement-regulatory protein and that complement activation has been implicated in aneurysm biology^44^. De novo CD46 engagement by VSMCs may reflect an attempt to regulate complement-mediated injury within the inflamed aneurysmal microenvironment. Collectively, these predicted changes identify VSMC-PVAT signaling axes that are concordant with established disease mechanisms and that may warrant direct experimental validation; we note, however, that these inferences derive from computational interaction prediction on combined datasets and require confirmation at the protein and functional level.

#### Strengths and Limitations

To the best of our knowledge, this is the most comprehensive single nuclear atlas of human cardiac and perivascular adipose depots created thus far, including LA, RCA and AA, and SAT samples across 10 donors, with all donors contributing samples from at least two depots and 6 donors contributing samples from all four depots. Because we sampled these depots from the same participants, we were able to make within-subject comparisons that control for inter-individual variability and isolate the effect of anatomic location. Our use of primary human tissue, together with an integrated analysis of cellular composition, transcriptional programs, and intercellular communication across health and disease states, we provide a framework for regional differences in adipose biology. Our study also has several limitations. First, the cell-cell interactions we report are computational predictions based on ligand-receptor co-expression, and require validation at the protein and functional level. Second, the disease-state analyses integrate our adipose data with external cardiomyocytes and VSMC datasets, introducing possible batch effects; we also used right ventricular cardiomyocytes as the closest available proxy for right coronary artery fat. Third, our participants were undergoing cardiac surgery and may not represent the general population. Fourth, with ten individuals and a low immune cell yield, we may be underpowered to detect inter-individual heterogeneity and to resolve immune subpopulations.

## Conclusion

Our data and analysis establish that human EAT and PVAT are not uniform depots but regionally specialized compartments whose cellular composition, transcriptional programs, and intercellular signaling are shaped by anatomic location and remodeled in the settling of AF and aortic aneurysm. These findings provide a framework for understanding how regional cardiac and vascular adipose tissue contribute to cardiovascular disease and identified specific signaling axes, particularly left atrial-enriched EPHA signaling and the disease-associated FN1, EGF, SLIT, NOTCH, and CD46 changes, as candidates for future mechanistic and therapeutic investigation.

## Acknowledgements

The authors thank the patients who agreed to participate in this research.

