## Supplementary Figures 1-5 for "Single-Nuclear RNA Sequencing Reveals Regional Specialization and Cellular Interactions in Epicardial and Perivascular Adipose Tissue"

**Supplementary data**

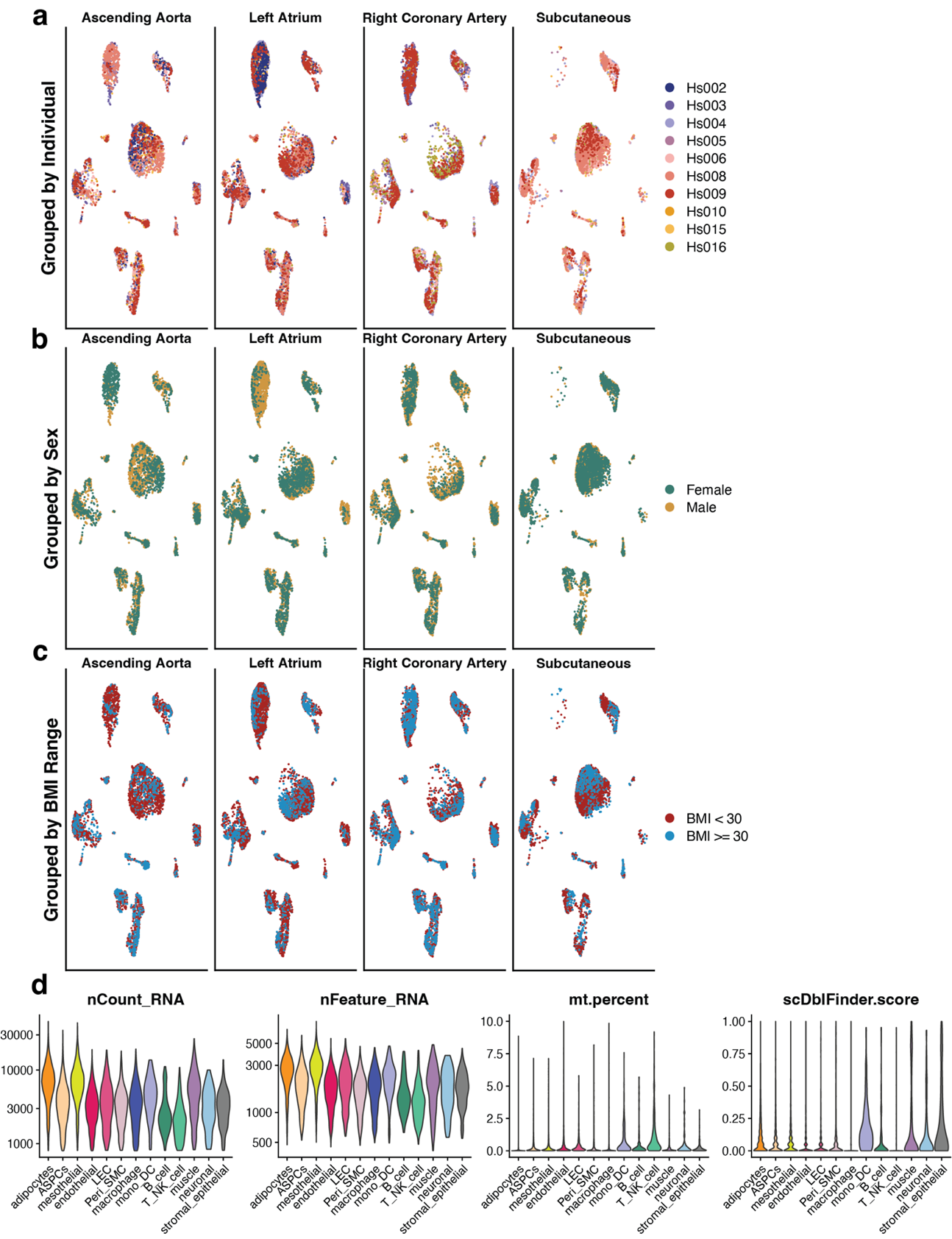

**Supplementary Figure 1. Distribution of the single nuclear dataset. a-c,** UMAP plots of integrated data from all depots, plotted by individual (**a**), sex (**b**), or BMI range (**c**). **d.** Quality control metrics for all clusters in the dataset.

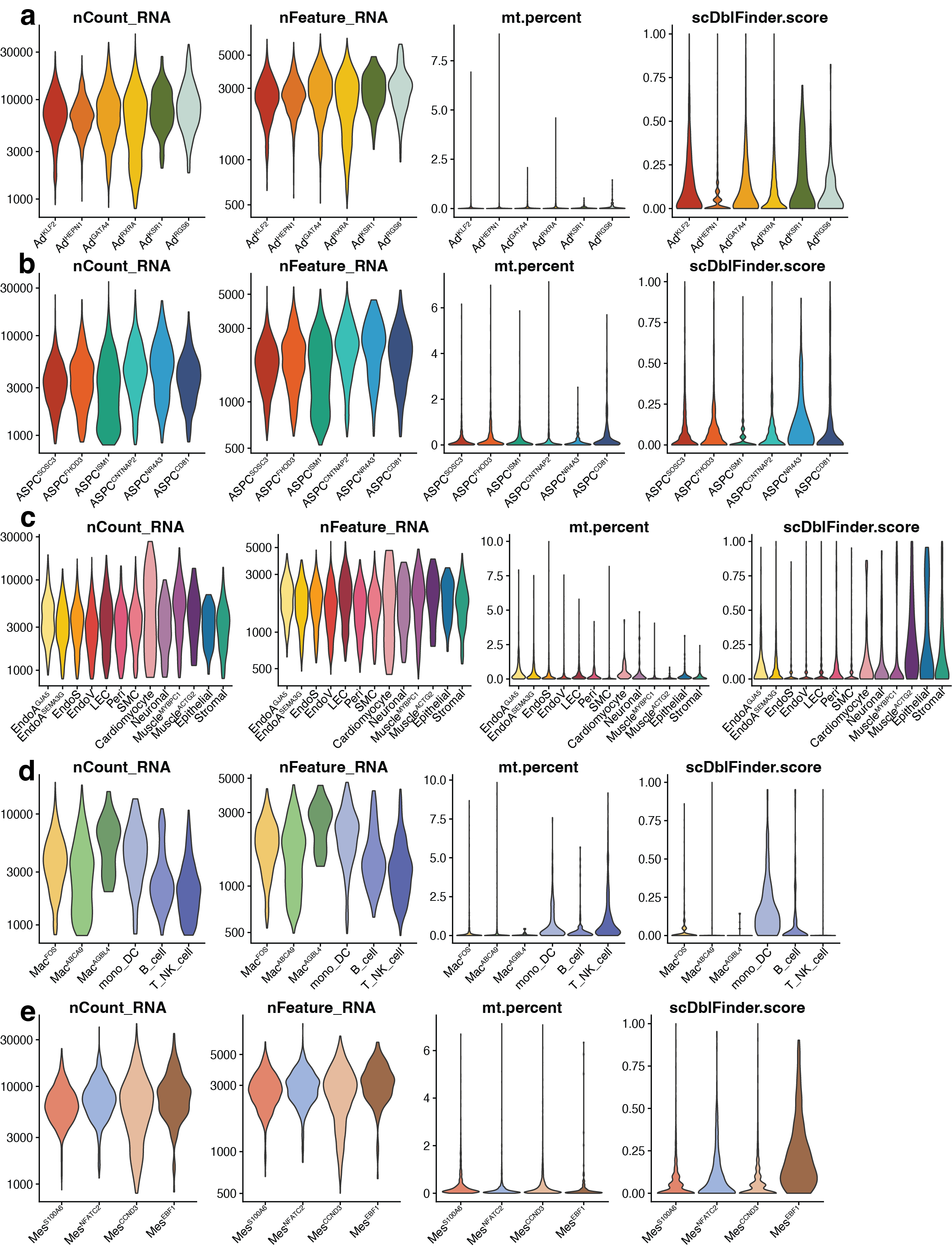

**Supplementary Figure 2. Quality control metrics for subclusters. a,** Quality control metrics for adipocyte subclusters. **b,** Quality control metrics for ASPC subclusters. **c,** Quality control metrics for vascular subclusters. **d,** Quality control metrics for immune subclusters. **e,** Quality control metrics for mesothelial subclusters.
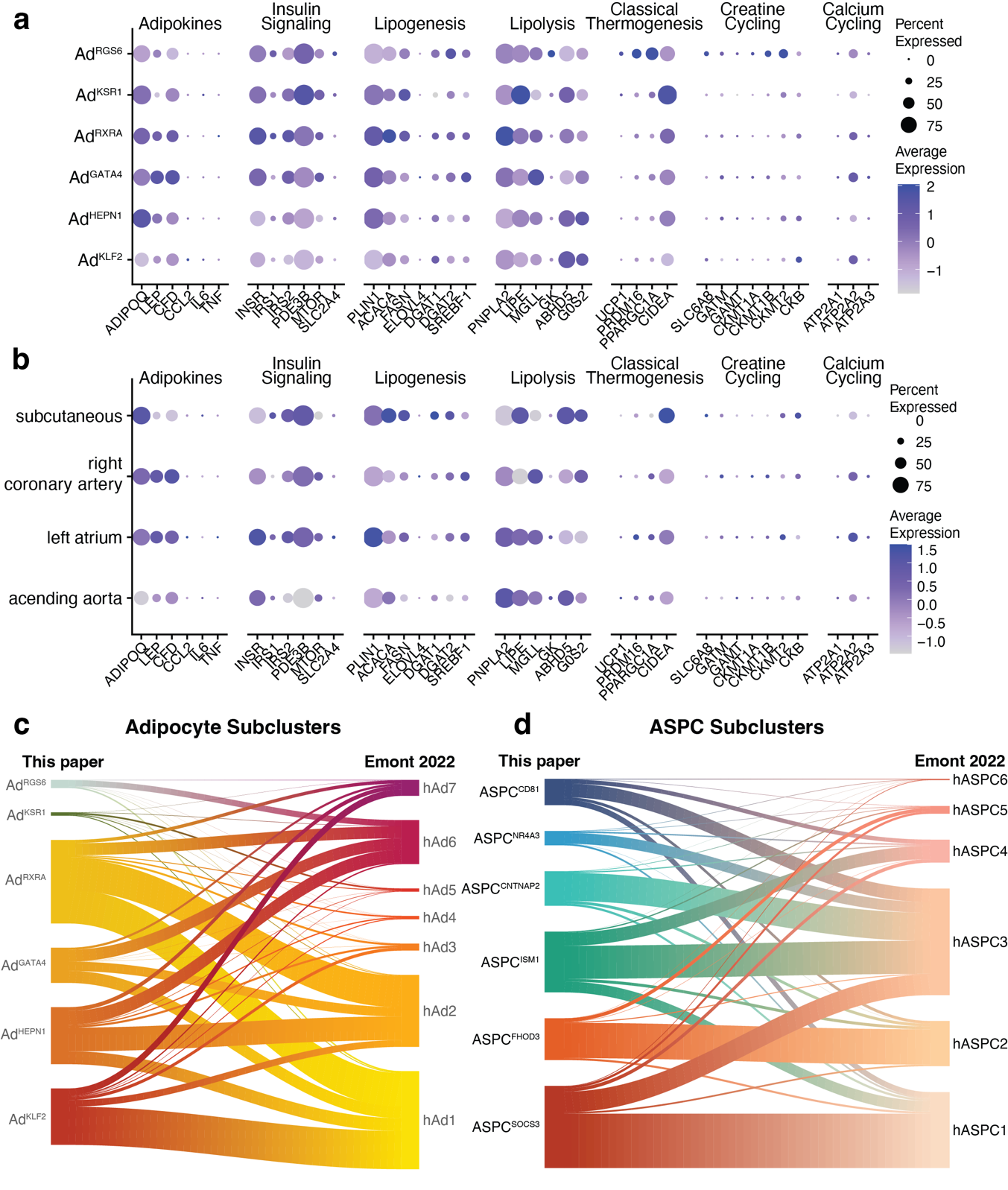

**Supplementary Figure 3. Further characterization of adipocyte and ASPC subclusters. a&b,** Genes associated with adipokines, insulin signaling, lipid handling, and thermogenesis plotted on adipocyte subclusters (**a**) or in all adipocytes from each depot (**b**).**c&d,** Riverplot comparing adipocyte (**c**) and ASPC (**d**) subclusters in this paper to those published in Emont 2022.

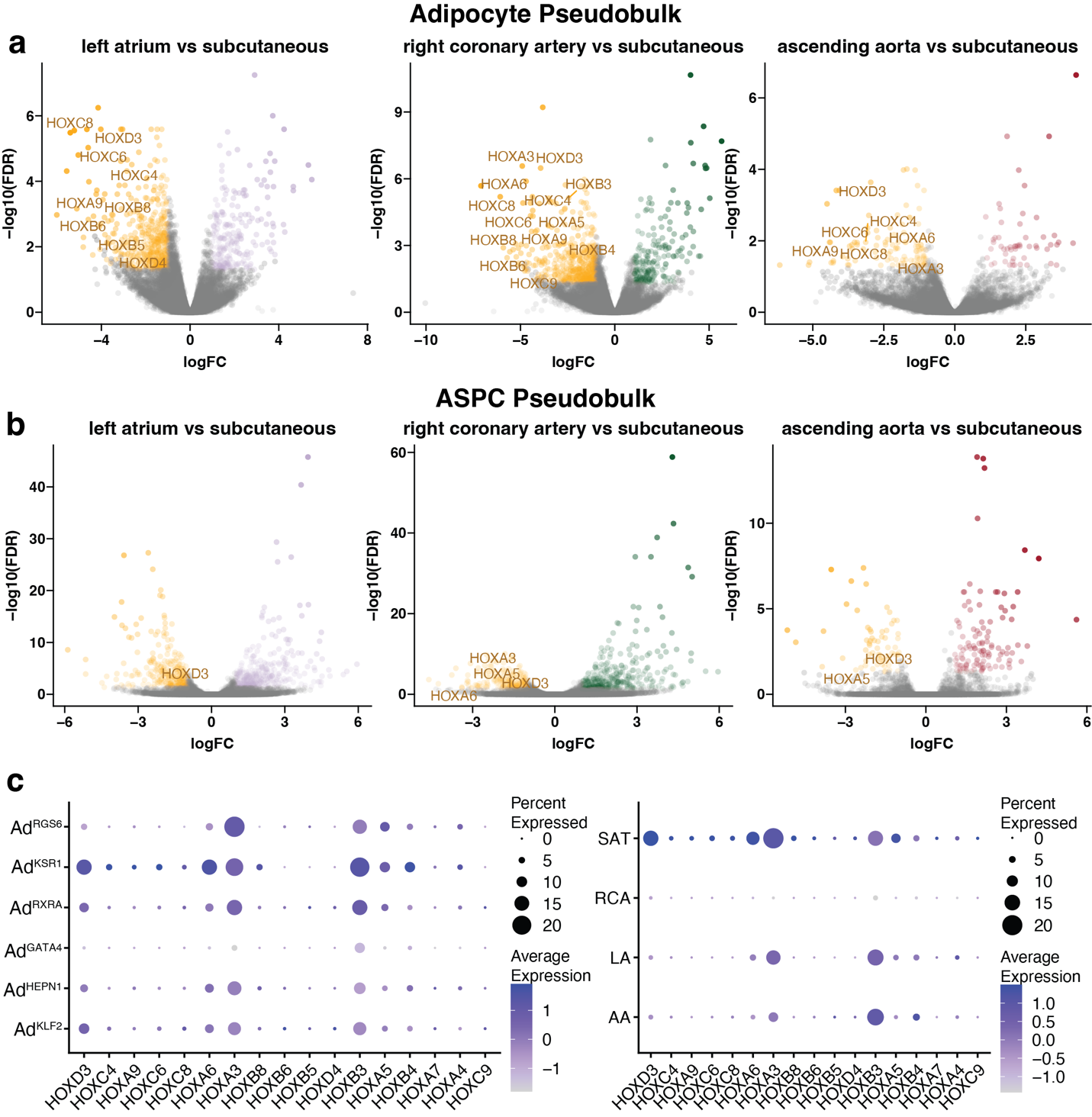

**Supplementary Figure 4. HOX gene expression across depot. a&b,** Volcano plots comparing gene expression in adipocytes (**a**) or ASPCs (**b**) from each epicardial depot compared to subcutaneous adipocytes with all significantly differential HOX genes highlighted. **c,** Dot plots showing the expression of HOX genes in adipocytes grouped by subcluster (**left**) or depot (**right**).

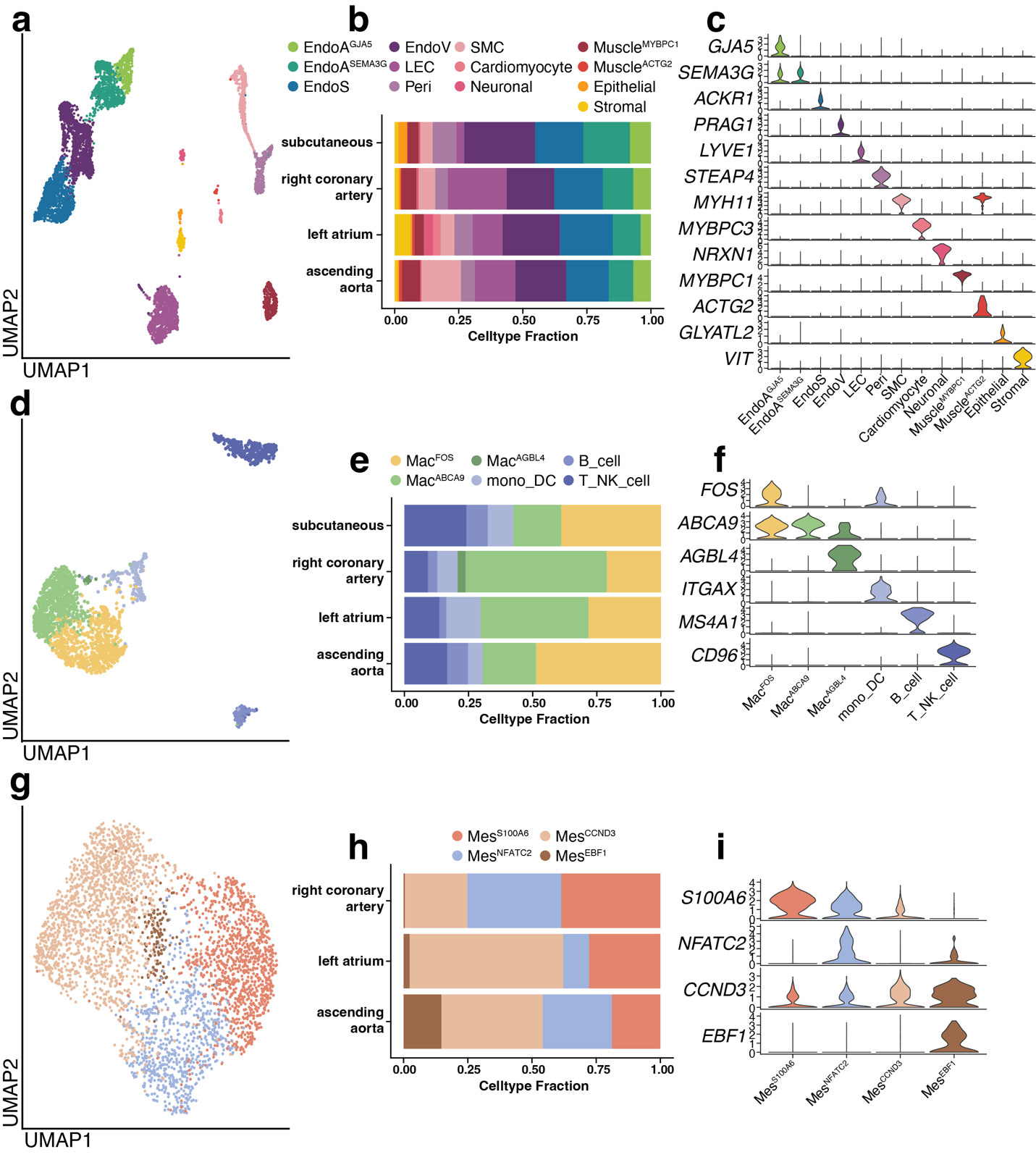

**Supplementary Figure 5. Subclustering of adipose tissue cell types. a,** UMAP plot of integrated vascular subclusters (n = 6288 cells). **b,** Cell type distribution of vascular subclusters across studied depot. **c,** Marker genes for vascular subclusters. d, UMAP plot of integrated immune subclusters (n = 2186 cells). **e,** Cell type distribution of immune subclusters across depot. **f,** Marker genes for immune subclusters. **g,** UMAP plot of integrated mesothelial subclusters (n = 3647 cells). **h,** Cell type distribution of mesothelial subclusters. **i,** Marker genes for mesothelial subclusters.
